# Optimizing the use of the Men-ACWY conjugated vaccine to control the developing meningococcal W disease outbreak in the Netherlands, a rapid analysis

**DOI:** 10.1101/494492

**Authors:** Albert Jan van Hoek, Mirjam Knol, Hester de Melker, Jacco Wallinga

**Author notes:** **Correspondence and requests for reprints to**: Albert Jan van Hoek, National Institute of Public Health and the Environment, P.O. Box 1, 3720 BA Bilthoven, The Netherlands.

## Abstract

**Background:** There is a developing outbreak of *Neisseria meningitidis* serotype W (MenW) in the Netherlands. In response, those aged 14 months and 14 years are vaccinated with the conjugated MenACWY vaccine. In the spring of 2018 we aimed to explore the impact of adding a one-off catch-up campaign targeting those aged 15-18 years on the transmission of MenW and the cost-effectiveness of such a campaign.

**Methods:** We estimated the growth rate of the MenW outbreak and quantified the impact of various targeted vaccination strategies on the reproductive number, and subsequently projected the future incidence with and without vaccination. Future cases were expressed in costs and QALYS and the incremental cost-effectiveness ratio was obtained.

**Results:** We estimate a reproductive number of around 1.4 (95%CI 1.2-1.7) over the period February 2016-February 2018. Adding the catch-up campaign reduces the reproductive number five years earlier than without a catch-up campaign, to a level around 1.2. The vaccination campaign, including the catch-up, will prevent around 100 cases per year in our base case scenario. Given the projected impact and realistic assumptions on costs and QALYs, adding the catch-up can be considered cost-effective using a threshold of €20,000 per QALY.

**Conclusion:** Adding the catch-up campaign targeting those aged 15-18 brings the impact of vaccination on reducing transmission five years forward and directly prevents a high-incidence age group from carriage and disease. Such a campaign can be considered cost-effective. Our study did underpin the decision to introduce a catch-up campaign in spring 2019. Furthermore, our applied method can be of interest for anyone solving vaccine allocation questions in a developing outbreak.

## Introduction

There is an ongoing *Neisseria meningitidis* serotype W (MenW) outbreak in the Netherlands due to a clonal complex 11 strain with 159 laboratory confirmed cases and 23 deaths between January 2015and February 2018 (1,2). As an outbreak response measure the conjugate MenC vaccine given at 14 months of age was replaced with the conjugate MenACWY vaccine in May 2018. In addition, in the autumn of 2018 a dose of the MenACWY vaccine was given to 14-year-old adolescents, with a continued vaccination of those who turn 14 in 2019 and 2020 (1). The Dutch Health Council will advise whether to continue this adolescent vaccination after 2020.

The aim of the vaccination programme is not only to protect those vaccinated against the development of disease, but also to reduce transmission and prevent disease and mortality in those not vaccinated, including those too young to receive the childhood dose and those very old.

In this paper we describe an analysis we performed in March and May 2018, based on the available data at that time, estimating the impact of the then planned programme on the transmission of MenW in the Netherlands, and exploring the additional impact of vaccinating those 15 to 18 years old in a one-off catch-up campaign. We aimed to provide evidence to inform the then pending decision whether or not to implement such a campaign.

To answer this vaccine allocation question we applied a state-of-the-art method, using limited amount of data, to quantify the impact of the vaccination programme on transmission. Subsequently we quantified the disease burden in costs and quality adjusted life years (QALY) and calculated incremental cost-effectiveness ratios (ICERs) for adding the catch-up campaign to the then current vaccination programme of a dose of Men-ACWY at 14 months and 14 years. Our answer informed the Outbreak Management Team meeting of the Netherlands in July 2018, and underpinned a decision to introduce the catch-up campaign, to be implemented in the first half of 2019.

## Methods

### Surveillance data

We used monthly-binned and aged-binned cumulative data from the Netherlands Reference laboratory for Bacterial Meningitis (NRLBM) from January 2015 to February 2018. Hence, the data are from the period before any vaccination programme was introduced. A detailed description of the surveillance system of meningococcal disease in the Netherlands is available in the recent publication by Knol *et al*. (2). We calculated the cumulative incidence in the following age groups: 0-1 years, 2-4 years, 5-13 years, 14-18 years, 19-24 years, 25-44 years, 64-79 years, 80+ years.

### Growth rate and reproductive number

We estimated the reproductive number (*R*_n_), the average number of secondary infections of an infected case, for a moving time-window of 25 months (*R*_n_ for January 2016 covers the period January 2015 until January 2017). For this period a growth rate (*r*) was obtained by Poisson regression of the cases by month using the number of months as an explanatory variable. We assume a carriage duration (*T*_c_) of 11 months (3) and are exponential distributed. Given this exponentially distribution, the generation interval will be exponential distributed too, which allowed us to calculate the reproductive number using formula 1 (4):

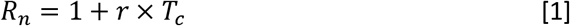

### Estimation impact of vaccination on the reproductive number

We estimated the reduction of R_n_ by immunizing specific age groups. When the reproductive number is reduced below one, the average number of secondary infections drops below the replacement threshold, and the transmission is likely to fade out. As no information on carriage and transmission of MenW is available, we were required to make several assumptions to interpret the clinical data. First, as the age distribution of cases is stable over time we assumed that the risk of disease given infection (case:carrier ratio) is also stable. Secondly, we assume that this case:carrier ratio is similar between the age of 5 until 79 years. This assumption is based on disease and carriage data for several meningococcal types, which showed clear elevated case:carrier ratios for the young and very old, but not for those age groups in between (5). For those younger (<5 years) and older age groups (80+ years) we thus assume that these groups contribute negligible to the transmission in carriage despite a, sometimes, high incidence. Thirdly, we assume that the proportion of the population which is susceptible to infection is similar between the age of 5 and 79 years. Fourth, we assume that infectious contacts are reciprocal (if John can infect Mary, Mary can also infect John). These four assumptions allow us to interpret the incidence by age as a force of infection (as in a hazard to become a carrier), by age. The change in the force of infection after vaccination can be estimated by applying the vaccine efficacy and vaccine coverage on the vaccinated age group. See Wallinga *et al*. PNAS 2010 for a full breakdown of the approach (6).

### Vaccine effectiveness and vaccination strategies

We assumed a vaccine effectiveness of 95% against disease with a slow, exponential, decline in protection after, such that it takes 20 years for 50% to loose protection. The vaccine efficacy against acquisition of MenW in carriage was set at 60% with 50% losing protection in 10 years (see (7) & appendix I). The assumed coverage is 95% for the dose at 14 months and 85% for adolescents.

We explored three different vaccination programmes which were relevant in the context of the Dutch situation. These are:

- vaccinating those aged 14 months annually
- vaccinating those aged 14 months annually + those aged 14 years annually
- vaccinating those aged 14 months annually + those aged 14 years annually + a one off catch-up of those aged 15 to 18 years old

For these three scenarios we estimated the reduction in R_n_ over time. As a sensitivity analysis we performed the same analysis for the previous outbreak of MenC in the Netherlands (data from January 1995 to May 2002 for selected type MenC:2a). In addition, we also explored vaccination at 6 months of age with a two-dose schedule.

### Projection future incidence without and with vaccination

Carriage (and transmission) of meningococcus does not result in lifelong protection against re-infection (5). Following this susceptible-infectious-susceptible (SIS) disease model, we project the future incidence by a logistical curve which shows an increase in the incidence of disease (see formula 2 & 3), and which stabilizes on a higher level. We use the estimated growth rate (*r*) and an assumption of the final incidence to parameterize this function, as although, we estimate the growth rate from the case data, we are not able to predict the future final incidence. Therefore, we explore the level of future incidence with a multiplier of three, as with this multiplier the incidence was similar to the height of the previous MenC outbreak before the introduction of a vaccination campaign (8). The age-specific incidence at time t0 (January 2018) is estimated from the age-specific incidence as observed in 2016 and 2017 by a least-squares approach. The indirect effect of the vaccination programme on the disease incidence in all age-groups (including those who are not vaccinated) is based on the annual estimated R_n_ as described in the previous paragraph. Vaccinated cohorts also enjoy a direct protection based on the vaccine efficacy (and coverage).

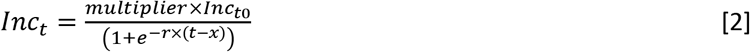

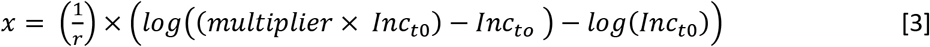

### Costs and QALY estimates

The estimation of costs and quality of life assumptions related to MenW disease are obtained from the last published cost-effectiveness analysis of the Men-ACWY vaccine in the Netherlands (9). The estimates of these costs and QALYs are reproduced in table 1 for clarity. As the costs in the previous publication were set for 2011, we inflated these to 2017 using the consumer price index (10).

**Table 1.** Incremental cost-effectiveness ratio of adding doses incrementally to only vaccinating those at 14 months of age, taking into account direct and indirect-effects of vaccination (Multiplier = 3).

| Target group | ICER (direct and indirect effects) |
| --- | --- |
| Only vaccinating at 14 months | € 34,556 (€ 25,941 - € 45,960) |
| + Adding those aged 14 years | € 26,985 (€ 20,129 - € 35,762) |
| + Adding those aged 15 to 18 | € 17,247 (€ 12,641 - € 23,204) |

### Mortality

The risk to die from an infection is based on the observed mortality during this outbreak. From the 159 identified cases between January 2015 and February 2018, 158 had information on death and23 died (15%) [data RIVM,(1)]. As there is no significant difference in mortality by age this percentage is applied irrespective of age. The 2018 life-table is used to estimate the (discounted) QALY adjusted life-expectancy, using background QALYs as collected by Versteegh *et al*. (11). In case of life-long sequelae the overall QALY adjusted life-expectancy is calculated using the same life-expectancy, but with a lower quality of life. The life-long costs are estimated using a similar approach.

### ICU use and duration of stay

The percentage of patients who suffer from a septic shock are based on the percentage of patients admitted to ICU during the historical MenC disease outbreak in the Netherlands which was also caused by clonal complex 11 (12). The duration of ICU admission and/or hospital stay are based on the same source.

### Vaccine and programme costs

We assume a vaccine price of €30 per MenACWY dose, and €10 per MenC dose. Hence, for the childhood MenACWY dose only €20 was included. The administration costs were set on €20 for the adolescent dose, and €0 for the replaced childhood dose. The additional costs to introduce the MenACWY programme among adolescents, thus the costs on top of purchasing the vaccine and the costs for administration, would be around €850,000 in the first year, €500,000 in the second year and only €4,0 00 in the third year and consecutive years [personal communication RIVM].

### Cost-effectiveness model

The costs and QALYs are discounted according to Dutch guidelines; costs by 4% and QALYs by 1.5% (13). The incremental cost-effectiveness ratio is calculated from a health care payer’s perspective. A threshold of €20,000 per QALY is used to categorize ICERs as being cost-effective, which is in line with common practice, although no formal threshold exists in the Netherlands. An overview of all costs and QALY assumptions is given in table S3-1 of the appendix.

## Results

### The reproductive number

From January 2015 until February 2018 there were 159 confirmed cases of MenW disease. The incidence continued to increase over time, with 10 and 9 reported cases in the first two months of 2018 respectively. Based on the observed growth the reproductive number declined from 2.3 (95% CI: 1.9-2.8) in the period January 2015-January 2017 to 1.4 (95% CI: 1.2-1.7) in the period February 2016-February 2018, see figure 1.

**Figure 1.**
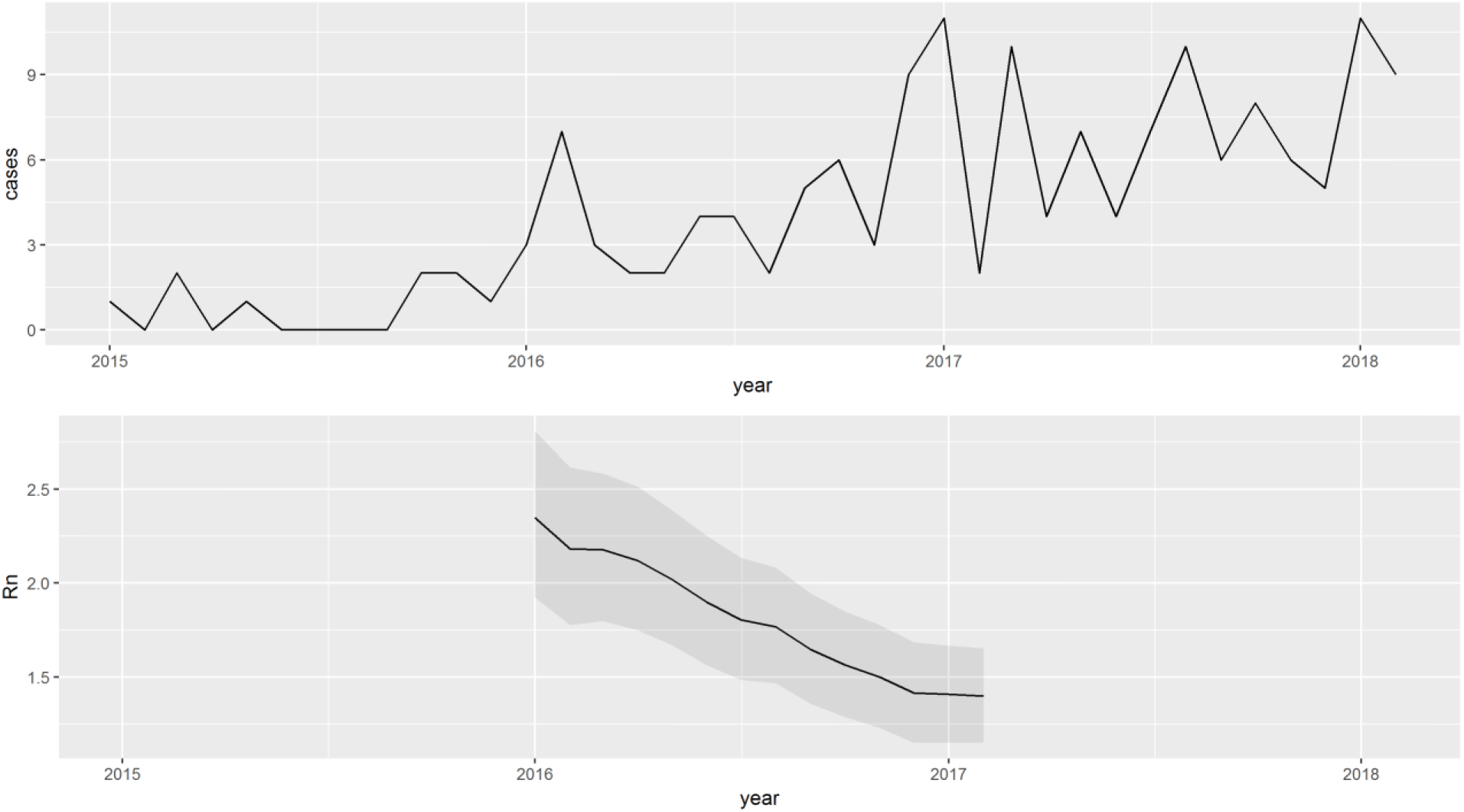
Overview of the cases per month for MenW in the period January 2015 until February 2018 (upper panel) as well as the estimated reproductive number (R_n_) over time (lower panel).

### Cumulative incidence by age

The cumulative incidence by age of MenW disease in the Netherlands is characterized by a relatively high incidence among those under 2 years (3.2 per 100,000) and over 80 years of age (2.9 per 100,000), see figure 2. The third largest incidence is among adolescents aged 14 to 18 years, in this group the incidence is 1.8 per 100,000. Although the incidence is very low in 2-13 year-olds 25-44 year olds, the incidence among those 45 years and over is not much lower compared to the 19-24 year-olds.

**Figure 2.**
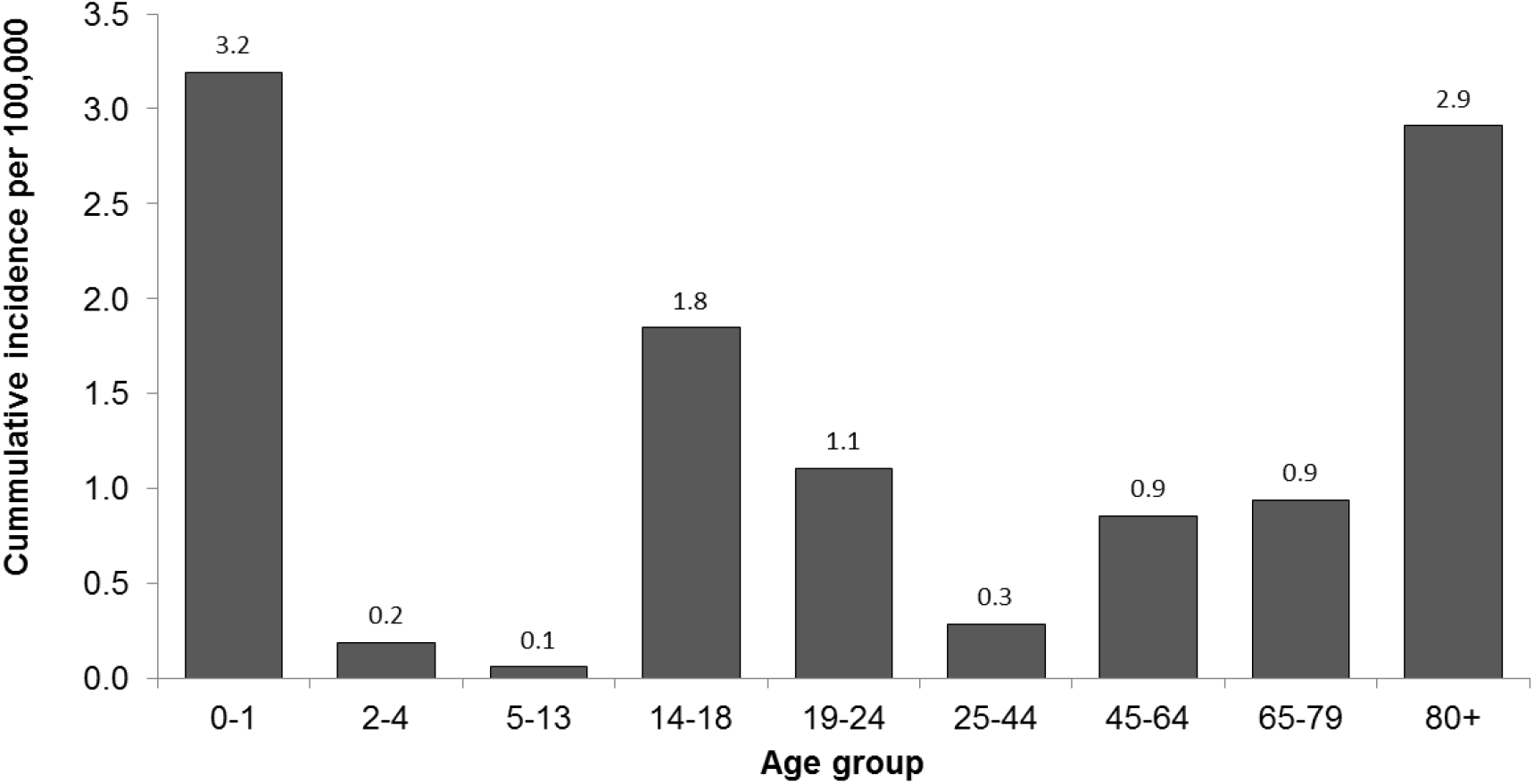
Cumulative incidence of MenW disease by age group, per 100,000

### Reduction of the reproductive number per 100,000 immunized, by age

Focusing only on those aged between 5 and 79 (because of the assumed similar case:carrier rate), the absolute reduction of the final R_n_ was estimated. For the MenW outbreak in the Netherlands this was 0.044 per 100,000 immunized (vaccinated and protected) in the age group 14-18 for the estimated final R_n_ of 1.4, see figure 3. For the previous MenC outbreak this was 0.168 per 100,000 immunized for an estimated R_n_ of 1.7, see appendix figure S2-3.

**Figure 3.**
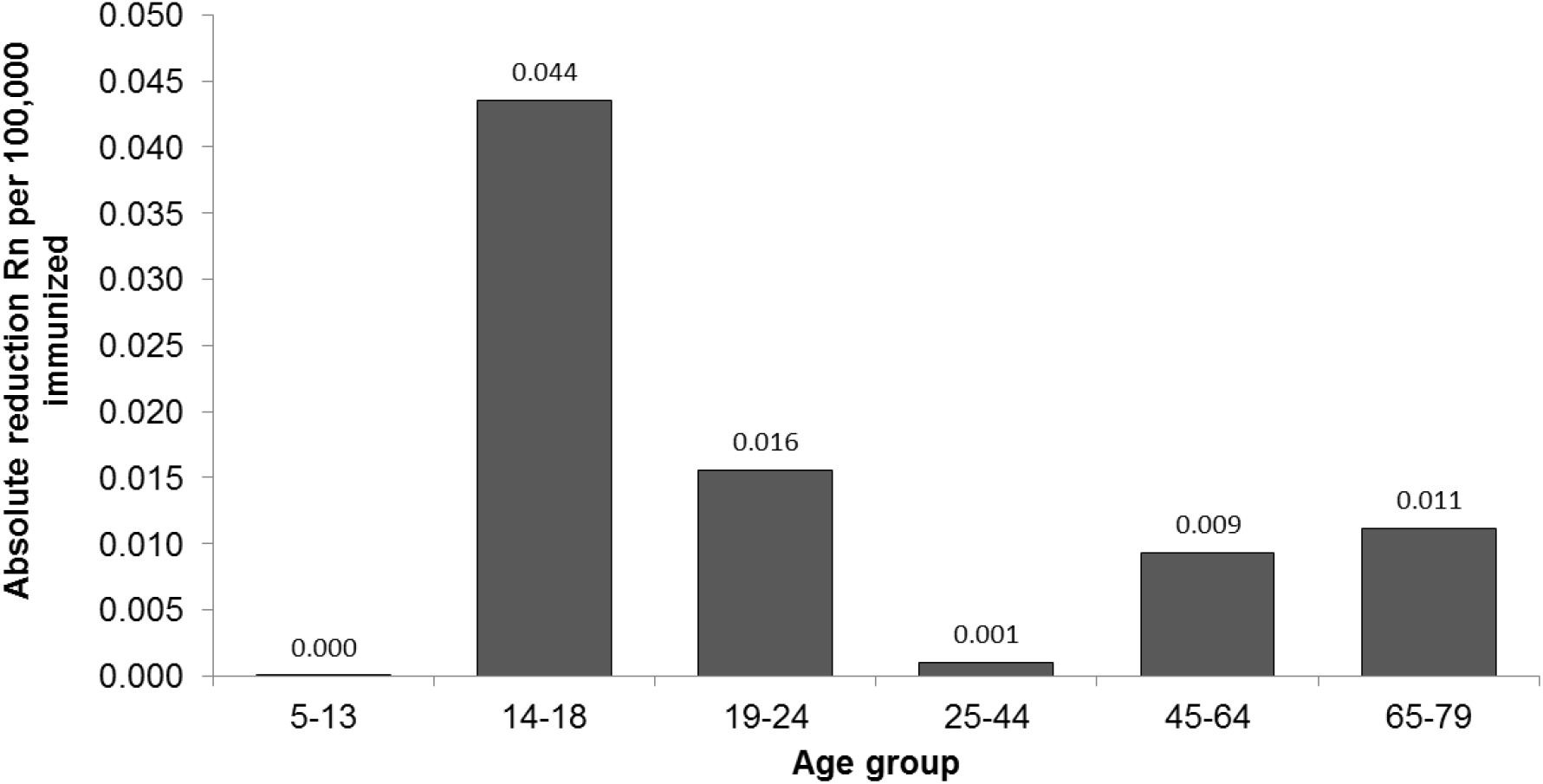
The estimated reduction of R_n_ per 100,000 immunized.

### Reduction of reproductive number over time, under different vaccination scenarios

Vaccination of 14 year-olds with a vaccine efficacy of 60% against carriage, an average duration of protection of 10 years and a coverage of 85%, on annual basis, will slowly decline R_n_ but the R_n_ is not likely to drop much below 1.2, see figure 4. Adding a catch-up among those aged 15 to 18 reduces R_n_ much faster, bringing the overall impact forward with 5 years. Although not shown, a longer duration of protection reduces the long-term equilibrium somewhat but does not lower the drop just after vaccination. In comparison, see appendix figure S2-4, the impact of vaccinating adolescents in the MenC outbreak has a much higher impact compared to MenW, as vaccination of adolescents is able to reduce the R_n_ just below one, even though the reproductive number before starting vaccination is higher.

**Figure 4.**
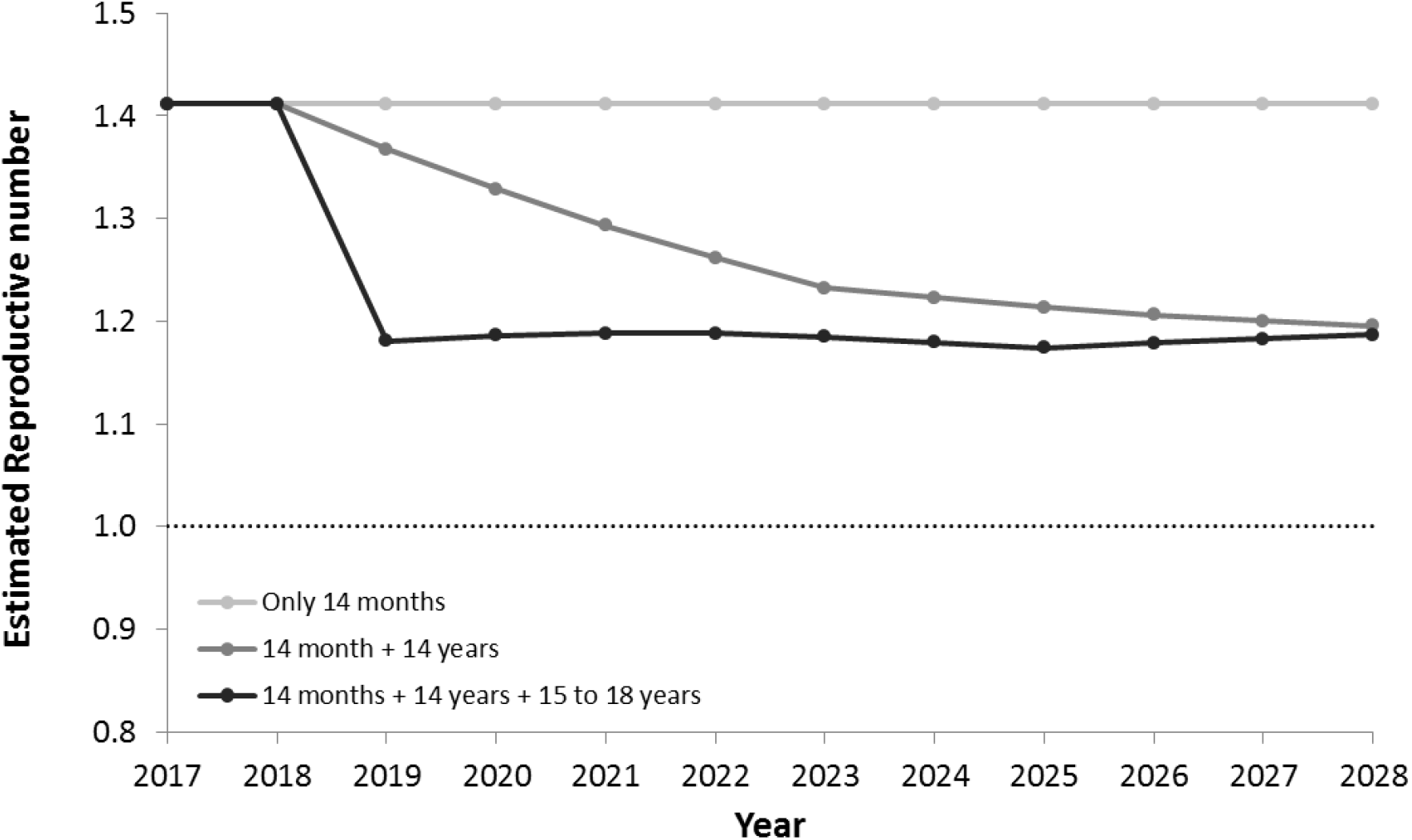
Estimated change in the reproductive number over time by the different vaccination programmes

### Projection future cases

Without any vaccination, and with a multiplier of three the projected number of cases will stabilize around 300 cases per year, see figure 5. With the current programme of a dose at 14 months and 14 years the annual number of cases will settle around 224. Adding the catch-up campaign will reduce this further to 201.

**Figure 5.**
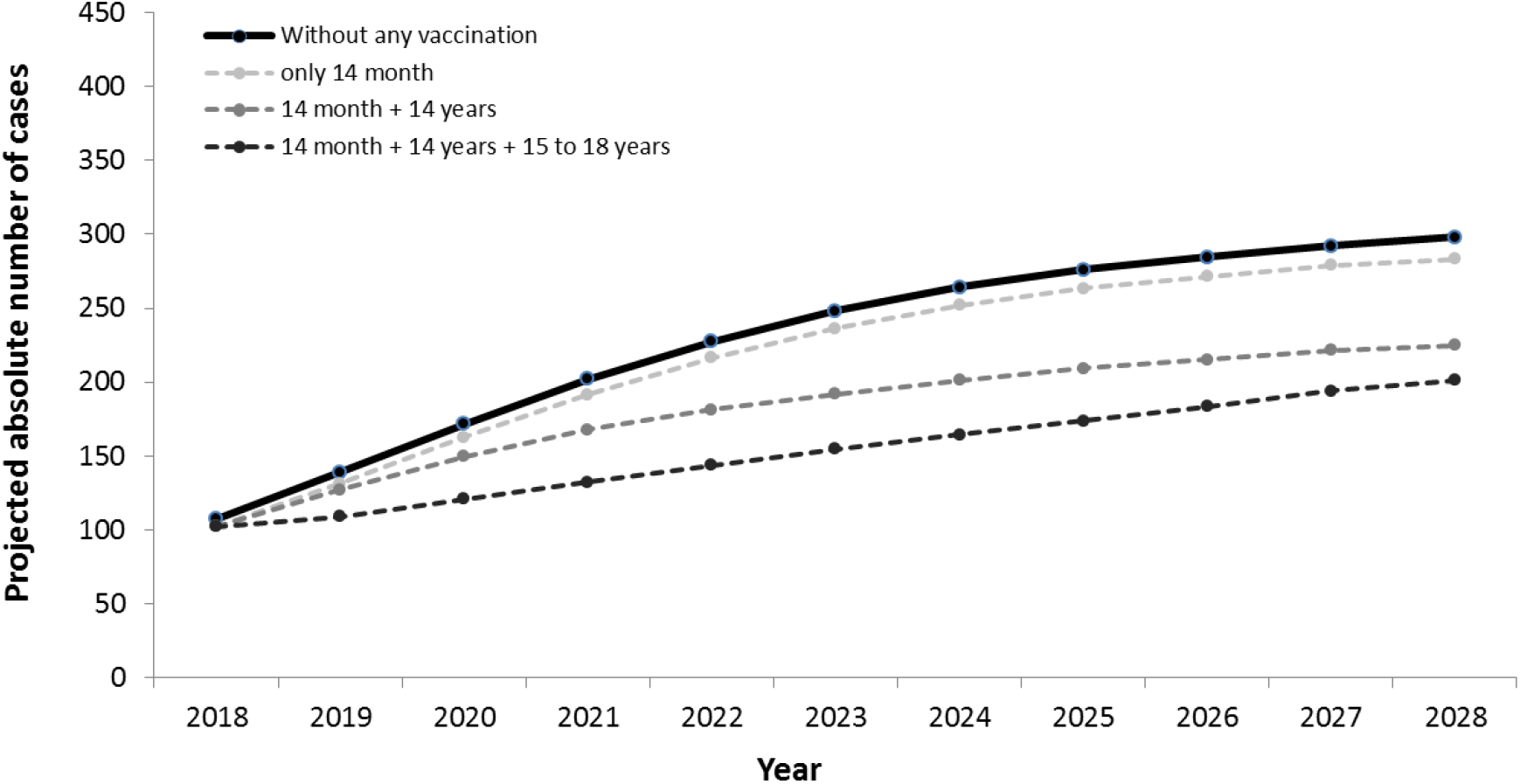
Projected absolute number of cases over the next 10 years, with different vaccination strategies

**Figure 6.**
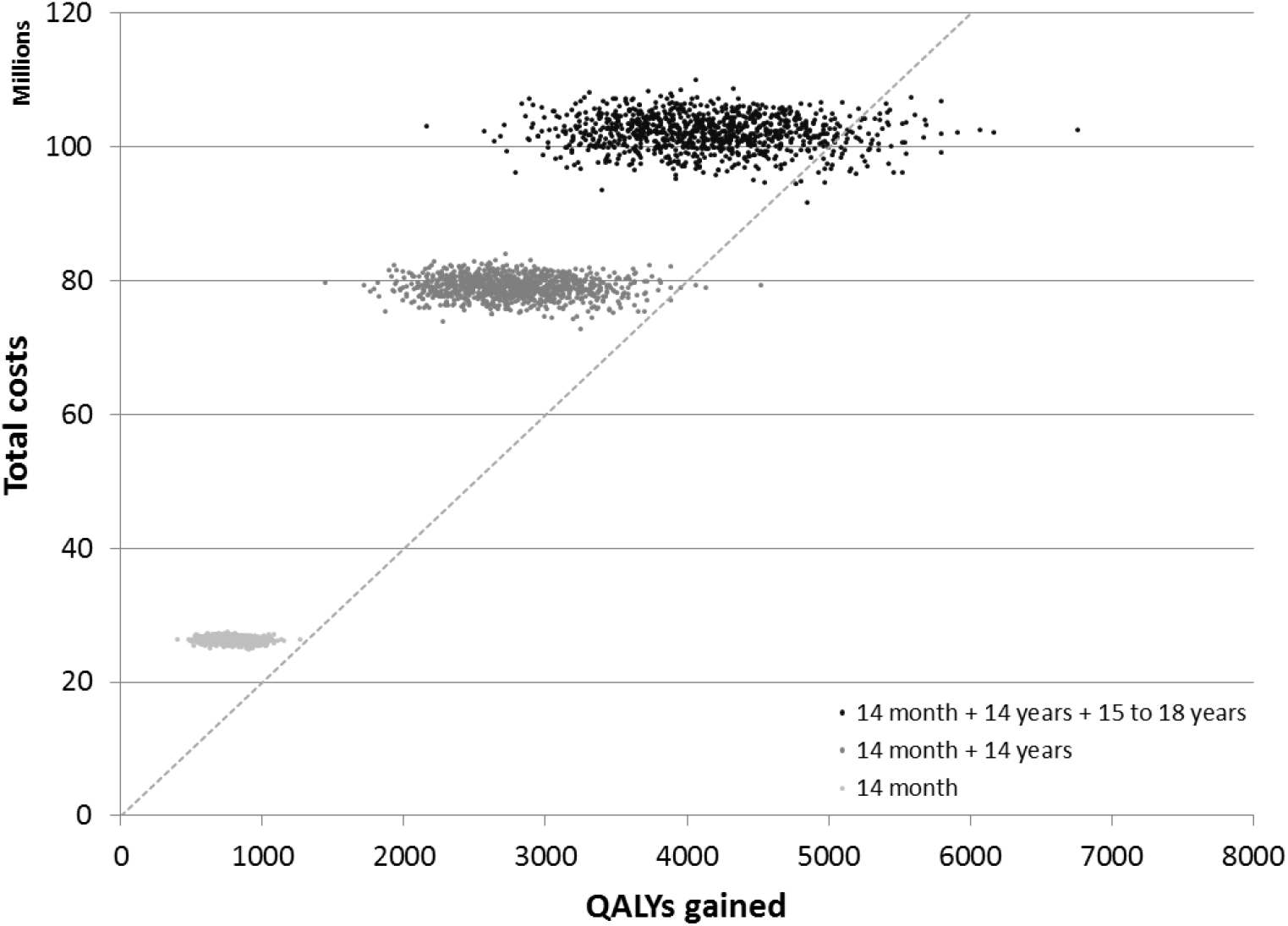
Cost-effectiveness planes showing the total costs and QALYs gained for the three different vaccination strategies (multiplier of 3).

### Cost-effectiveness

Without including the indirect effects, and only looking at individual programmatic options, vaccinating children at 14 months is most cost-effective, see table S3-2, as they require one dose to prevent the highest incidence of disease and is less expensive to implement. All CEAs remain however below €50,000 per QALY except vaccination at 6 months which requires two doses and is therefore much less cost-effective, see table S3-2 in the appendix. The most relevant comparison to address our question is however the incremental cost-effectiveness ratio of adding the catch-up campaign targeting those 15-18 years old to the already planned programme targeting 14 months and 14 years old and including both direct and indirect effects. In this case the ICER is € 17,247 (95% Credibility Interval (CrI): €12,641 - €23,204), see table 1. Therefore, adding this campaign can be considered cost-effective, although the confidence interval does include the €20,000 threshold.

### Sensitivity analysis

The cost-effectiveness of the different strategies is strongly dependent on factors such as the exact growth rate, the protection of the vaccine against carriage, the duration of carriage, and the multiplier. When the future incidence is higher, using a multiplier of four, the planned programme is cost-effective at € 19,028 (95% CrI: € 14,074 - € 25,280), however the addition of a catch-up campaign will still be even more cost-effective with an ICER of € 9,624 (95% CrI: € 6,722 - € 13,163), see table S3-4 in the appendix. Thus, adding the catch-up campaign is cost-effective as soon as the total number of MenW cases is expected to rise (without vaccination) until something close to 300 cases per year (multiplier of 3). An uptake of 85% in the catch-up is high, but with a lower uptake of 65% the ICER increases only marginally to €17,616 (95% CrI: € 12,917-€ 23,676).

The estimated growth rate and R_n_ are influential for the outcome. When the lowest end of the confidence interval of the estimated R_n_ is used (1.2) the growth of the outbreak is slower and already better to control with only vaccinating the 14 year olds. Due to this, using a 10 year time horizon, it is less cost-effective to implement any vaccination campaign, and adding a catch-up is slightly less cost-effective with an ICER of € 22,227 (95% CrI: € 16,411 - € 29,647). With the highest end of the R_n_ (1.7) the growth is much faster, but more difficult to control. The catch-up campaign does however lower the disease burden and is therefore still cost-effective, with an ICER of € 18,633 (95% CrI: € 13,376 - € 24,845).

## Discussion

Our analysis showed a growing outbreak with a reproductive number of around 1.4. The age distribution of confirmed cases of MenW disease suggested an important role of adolescents in the transmission. Adolescents are therefore the best target for a catch-up vaccination campaign with the conjugated MenACWY vaccine. Adding the catch-up campaign reduces the reproductive number five years earlier in time compared to not adding the catch-up, and reduces the expected cases. A delay in the implementation of the MenW vaccination programme will mean that the incidence will stabilize at a higher level, with more cases and deaths per year. Therefore, the sooner an intervention is widely implemented, the better. Given the projected impact and realistic assumptions on costs and QALYs, adding the catch-up can be considered (marginally) cost-effective using a threshold of €20,000 per QALY.

In addition, the age distribution of cases for MenW is different compared to the most recent MenC outbreak in the Netherlands. Applying the same assumptions for both diseases suggested a smaller effect of vaccination on the transmission of MenW compared to the impact on the MenC outbreak in 2002. Therefore, the experiences with the catch-up campaign for MenC is not necessary a blue-print for the expected impact of the catch-up campaign against MenW.

Given the uncertainties and limitations, and the very limited amount of data and knowledge about MenW, we would like to emphasize that our results should be treated as order-of-magnitude calculations. The duration of carriage for MenW is unknown, and we relied on an estimate of the duration which is not serotype specific and based on carriage among children aged 3-14 years (3). A shorter duration would lower our estimate of R_n_, and make it easier to control the outbreak, a longer duration would increase R_n_. Furthermore, nothing is known about the age specific carriage of MenW, and hence nothing can be said about the case:carrier ratio of MenW disease. For the carriage of other serogroups, there is a peak at age 19 years (14), however it is unclear if this is the case for MenW as well. Our assumption that the case:carrier rate is similar among the wide range of age groups needs confirmation, but our approach can be seen as conservative, as we lower the impact of vaccinating adolescents by including the possible transmission between adults and older adults. In our analysis we assumed an efficacy of 60% against acquisition of carriage. The only existing trial exploring the impact of MenACWY vaccine against carriage presents the odds of carriage after vaccination measured at 2, 4, 6, and 12 months and analyzed combined suggesting an efficacy of 36.2% (CI 95% 15.6-51.7) against carriage of serogroups CWY (7), which is therefore not the protection against acquisition of carriage. Given the long duration of carriage, the efficacy against acquisition is higher compared to this percentage. We therefore believe that our assumed 60% is in line with existing trial data. Nevertheless, there is conflicting evidence of carriage after vaccination (15,16), and therefore careful monitoring of indirect protection caused by MenACWY vaccination seems paramount.

To our knowledge, this is the first paper describing the application of our approach to quantify the reduction of the reproductive number and project the future incidence on a developing outbreak and combining it with a cost-effectiveness analysis. Given the afore mentioned limitations and uncertainties we expect that analyzing the same data with more complex methods, such as fitting a dynamic transmission model would lead to the same conclusions. This is important, as the method we applied can be performed in standard spreadsheet software and be implemented in a short time frame.

In conclusion, given an expected high efficacy against the prevention of disease we predict a strong reduction in disease among those vaccinated, but the indirect protection towards those not vaccinated might be slow to pick up, especially when there is no catch-up campaign, or where the coverage is low. Adding a catch-up campaign to the planned vaccination programme will halt the outbreak earlier and therefore prevent extra cases and deaths in those vaccinated, but also in those not vaccinated. And earlier implementation is better. Under the realistic future scenario of more than 200 cases per year without vaccination, adding a catch-up is a better strategy compared to the current implemented programme. Given the difference in age distribution of cases we should be cautious in using the previous experience with the successful vaccination campaign against MenC as indicative for the impact of an adolescent vaccination campaign against MenW in the Netherlands.

## Supporting information

