## Supplementary material for "Optimizing the use of the Men-ACWY conjugated vaccine to control the developing meningococcal W disease outbreak in the Netherlands, a rapid analysis"

**Appendix I**

### Vaccine efficacy against acquisition and vaccine efficacy against carriage

The vaccine efficacy of the conjugated Men-ACWY against carriage end-points has been measured in a clinical trial (1). The main end-point in the publication of this trial is the vaccine efficacy against carriage, as measured as 1-odds ratio of carriage in month 2, 4, 6 and 12 after vaccination. This measure differs from the vaccine efficacy as applied in our analysis, which is the vaccine efficacy against acquisition. The vaccine efficacy against acquisition is the reduction in acquiring Men W. There is a relation between the VE against carriage and the VE against acquisition, which depends on the duration of carriage. To have an understanding of which VE against acquisition might correspond to published VE of 34% against carriage, we simulated the carriage in 987 recipients of placebo and 998 recipients of the vaccine and a carriage duration of 11, 9 or 6 months. Assuming a constant acquisition rate, the acquisition rate was fitted to the observed 11% carriage (measured at month 2, 4, 6 and 12) in the placebo arm. In figure 1 we plotted the relation between the vaccine efficacy against carriage (% reduction in the carriage rate) and the vaccine efficacy against carriage based on the measured carriage at 2, 4, 6 and 12 months (1-Odds ratio) post vaccination. This reveals that the VE against acquisition is higher than the observed VE against carriage. Thus, even though the study observes a VE against carriage of 34% the corresponding VE against acquisition is higher. Under our assumption of a constant acquisition rate the vaccine efficacy of 34% against carriage corresponds to an efficacy against acquisition of 68% (11 months), 60% (9 months) and 49% (6 months). Even when the vaccine efficacy against acquisition is 100%, the maximum observed VE against carriage would have been 50% (11 months), 56% (9 months) and 69% (6 months). We applied a VE of 60% against acquisition in our trial, which we believe is in line with the results of this trial.

**Figure S1-1** Simulated relation between the VE against acquisition (% reduction of the acquisition rate) and the corresponding observed VE against carriage (1- Odds), as measured at month 2, 4, 6 and 12, for three different carriage durations (11, 9 & 6 months).

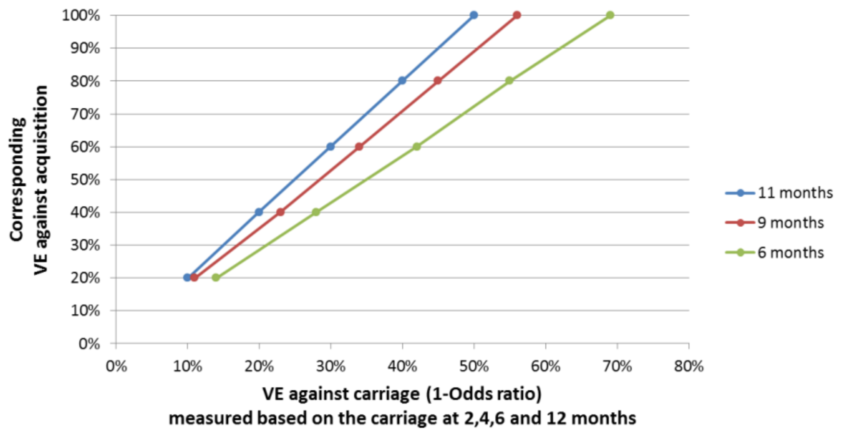

**Appendix II**

**The comparison between the estimated impact on the MenW outbreak and the MenC outbreak**

*Reproductive number outbreak MenC*

For the MenC outbreak there were 620 reported cases over the period January 1995 until May-2002. The reproductive number was constantly and significantly above 1 from the period Feb-1999 to Feb-2001 onwards. With an estimated Rn of 1.7 (95% CI: 1.6-1.9) in May 2002, see figure S2-1.

Figure S2-1. Overview of the case per month for MenC in the period January 1995 until May 2002 (upper panel) as well as the estimated reproductive number (Rn) over time (lower panel).

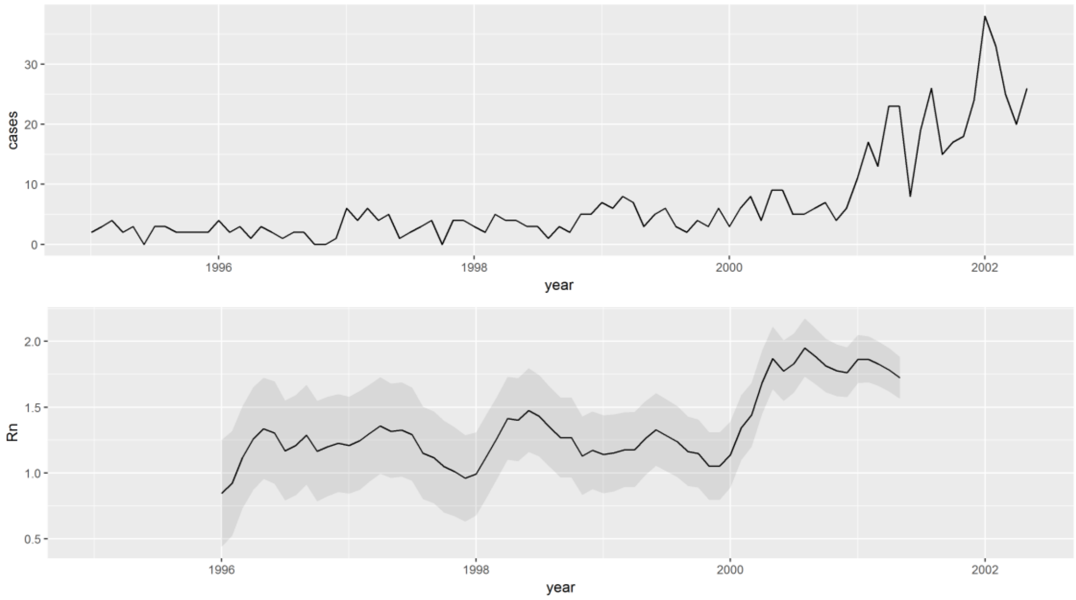

*Cumulative incidence by age*

The outbreak of MenC in the Netherlands was further developed compared to MenW, resulting in a higher cumulative incidence. However, importantly the age distribution of cases is very different, with a concentration of the disease burden among those younger than 25. With a much higher disease burden among 2 to 13-year old and a relatively low incidence among older age groups. Even though the absolute incidence among 80+ year old is similar compared to MenW, this age group is much less affected compared to the outbreak of MenW, see figure S2-2.

Figure S2-2 comparison of the cumulative incidence by age for MenW (2015-2018) and MenC (1995-2002).

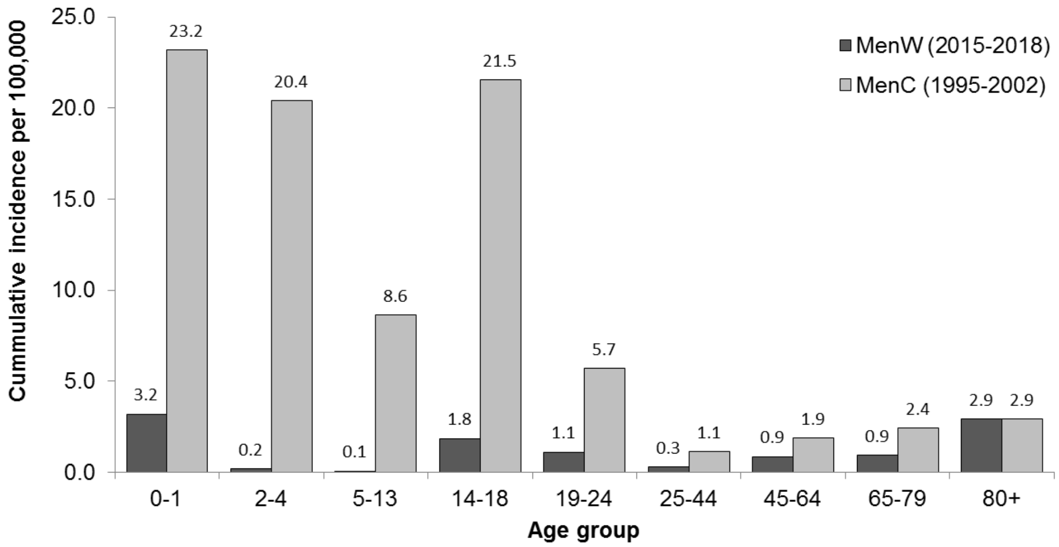

*Impact on the reproductive number*

For MenC the impact is very much concentrated in the age group 14-18, with an absolute reduction of Rn by 0.17 per 100,000 vaccinated linked to the final Rn of 1.7, with a very low impact of vaccinating older age groups, see figure S2-3.

Figure S2-3 Comparison of the estimated absolute reduction of the estimated reproductive number (1.4 MenW and 1.7 MenC) by immunizing 100,000 persons in each age group.

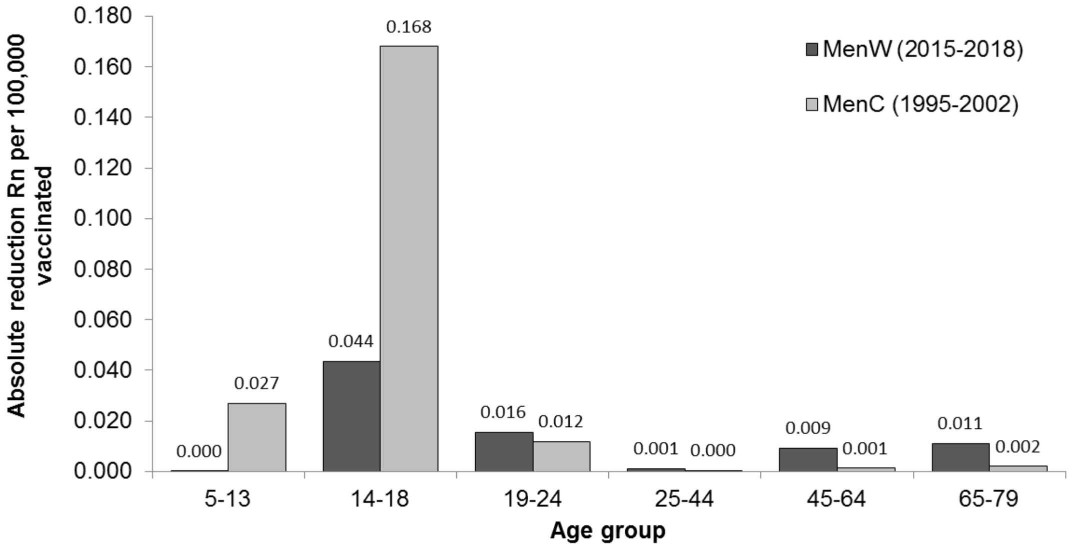

Figure S2-4 Impact of vaccination on the reduction of the reproductive number

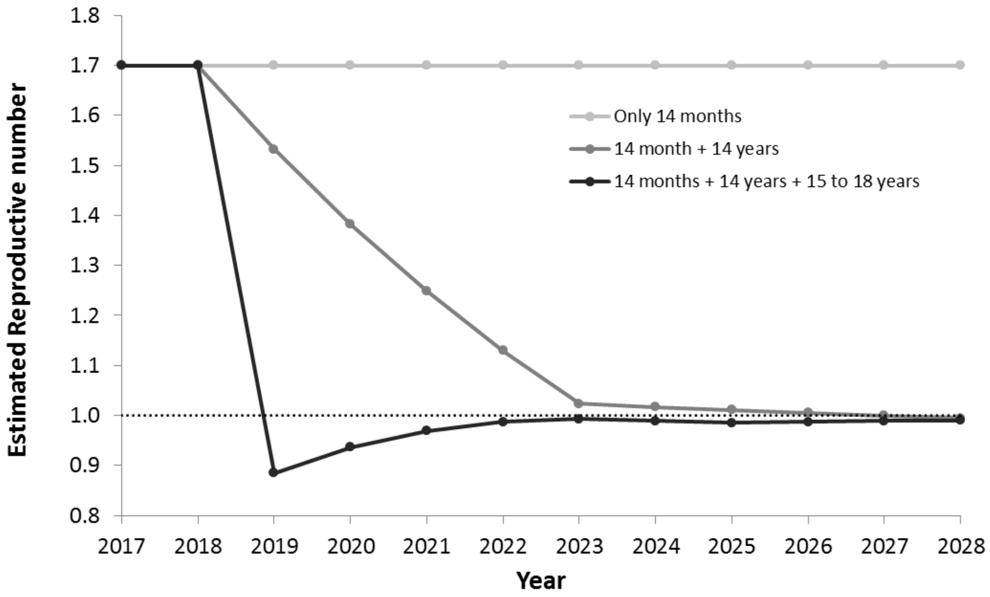

**Appendix III**

The assumptions and extra results for the cost-effectiveness analysis.

Table S3-1 Costs and QALY assumptions as used in the analysis

| **Parameter** | **Point estimate** | **Distribution** | **Reference** |
|---|---|---|---|
| Vaccine efficacy invasive disease (%) | 95% | | assumed |
| Waning of protection efficacy invasive disease | Over 20 years 50% loses protection | | assumed |
| Vaccine efficacy acquisition carriage (%) | 60% | | assumed |
| Waning of protection efficacy acquisition carriage | Over 10 years 50% loses protection | | assumed |
| Vaccination coverage (14 month) | 95% | | assumed |
| Vaccination coverage (adolescents) | 85% | | assumed |
| Percentage of Men W disease without septic shock | 46% | Beta(60;70) | (2) |
| Case fatality | 16% | Beta(29;151) | RIVM |
| *Sequelae (%)* | | | |
| Scars | 3.9% | Beta(20;495) | (3) |
| Amputations | 0.8% | Beta(4;511) | (3) |
| Neurological sequelae | 6.9% | Beta(127;1707) | (3) |
| *Price vaccine* | | | |
| Vaccine dose Men ACWY | € 30 | | assumed |
| Administration costs | € 10 | | assumed |
| Administration costs 14 months | € 0 | | assumed |
| Administration costs adolescents | € 20 | | assumed |
| *Extra costs adolescents campaign* | | | |
| First year | € 850,000 | | RIVM |
| Second year | € 500,000 | | RIVM |
| Third and subsequent years | € 4,000 | | RIVM |
| *Medical costs (2017 price level)* | | | |
| GP visit | € 31 | | (3) |
| Microbiological diagnostics and MRI | € 331 | Gamma(1;331) | (3) |
| Full course parenteral antibiotic treatment | € 322 | Gamma(1;322) | (3) |
| Inpatient day (general ward) | € 502 | | (3) |
| Inpatient day (intensive care unit) | € 2,455 | | (3) |
| Extra medical assistance with septic shock | € 2,039 | Gamma(1;2,039) | (3) |
| Treatment scars | € 549 | Gamma(1;549) | (3) |
| Treatment for amputations | € 1,766 | Gamma(1;1,766) | (3) |
| Institutional care (annual costs) | € 97,692 | Gamma(1;97,692) | (3) |
| Special education | € 5,100 - € 14,984 | Age specific | (3) |

| | | | |
|---|---|---|---|
| Pediatrician follow-up | € 132 | Gamma(1;132) | (3) |
| Public health follow-up | € 67 | Gamma(1;67) | (3) |
| *Duration of hospitalization* | | | |
| No septic shock; general ward | 10 | Gamma(1;10) | (2) |
| Septic shock; general ward | 10 | Gamma(1;10) | (2) |
| Septic shock; intensive care | 4 | Gamma(1;5) | (2) |
| Treatment scars | 2 | Gamma(1;2) | (3) |
| Treatment amputations | 8 | Gamma(1;8) | (3) |
| *Drop in quality of life (QALY)* | | | |
| Amputations or scars | 0.17 | | (3) |
| Neurological sequelae | 0.25 | | (3) |
| Average quality of life general population | Age dependent | | (3) |
| *Discount rate* | | | |
| Costs | 4% | | (4) |
| Health effects | 1.5% | | (4) |

Table S3-2 Cost-effectiveness ratio of each individual programme, taking into account only the direct effects or both the direct and indirect effects.

| Target group | ICER (direct effects only) | ICER (direct and indirect effects) |
|---|---|---|
| Only vaccinating at 6 months | € 89,773 (€ 67,489 - € 119,502) | Same |
| Only vaccinating at 14 months | € 34,556 (€ 25,941 - € 45,960) | Same |
| Only vaccinating at 14 years | € 39,769 (€ 29,958 - € 52,847) | € 26,092 (€ 19,435 - € 34,583) |
| Only vaccinating those aged 15 to 18 in a one off catch-up campaign | € 44,540 (€ 33,573 - € 59,194) | € 18,733 (€ 13,805- € 24,988) |

Table S3-3 Cost-effectiveness ratio of the incremental programme compared to doing nothing – thus the cost-effectiveness ratio linked to the clouds in figure 6 in the main paper.

| Target group | ICER (direct and indirect effects) |
|---|---|
| Only vaccinating at 14 months | € 34,556 (€ 25,941 - € 45,960) |
| + Adding those aged 14 years | € 29,101 (€ 21,828 - € 38,540) |
| + Adding those aged 15 to 18 | € 25,187 (€ 18,729 - € 33,406) |

Table S3-4 Sensitivity analysis

| Scenario | ICER (adding catch-up programme to current programme of vaccinating those aged 14 month & 14 years) |
|---|---|
| VE acquisition similar as VE against invasive disease (=95%) | € 7,836 (€ 5,320 - € 10,849) |
| Coverage catch-up 65% instead of 85% | €17,616 (€ 12,917- € 23,676) |
| Carriage duration is 9 months | € 16,955 (€ 12,303 - € 22,931) |
| Multiplier = 4 | € 9,624 (€ 6,722 - € 13,163) |
| Lower CI growth rate (R0 = 1.2) | € 22,227 (€ 16,411 - € 29,647) |
| Upper CI growth rate (R0=1.7) | € 18,633 (€ 13,376 - € 24,845) |

References appendix:

1. Read RC, Baxter D, Chadwick DR, Faust SN, Finn A, Gordon SB, et al. Effect of a quadrivalent meningococcal ACWY glycoconjugate or a serogroup B meningococcal vaccine on meningococcal carriage: An observer-blind, phase 3 randomised clinical trial. Lancet [Internet]. 2014;384(9960):2123–31. Available from: http://dx.doi.org/10.1016/S0140-6736(14)60842-4

2. Stoof SP, Rodenburg GD, Knol MJ, Rümke LW, Bovenkerk S, Berbers GAM, et al. Disease Burden of Invasive Meningococcal Disease in the Netherlands between June 1999 and June 2011: A Subjective Role for Serogroup and Clonal Complex. Clin Infect Dis. 2015;61(8):1281–92.

3. Hepkema H, Pouwels KB, van der Ende A, Westra TA, Postma MJ. Meningococcal Serogroup A, C, W135 and Y Conjugated Vaccine: A Cost-Effectiveness Analysis in the Netherlands. PLoS One. 2013;8(5):1–11.

4. Zorginstituut Nederland. Guideline for economic evaluations in healthcare. 2016;1–45. Available from: https://english.zorginstituutnederland.nl/publications/reports/2016/06/16/guideline-for-economic-evaluations-in-healthcare
